# Characterization of chondroitinase-induced lumbar intervertebral disc degeneration in a sheep model intended for assessing biomaterials

**DOI:** 10.1101/2020.05.25.115667

**Authors:** Ryan Borem, Joshua Walters, Allison Madeline, Lee Madeline, Sanjitpal Gill, Jeremiah Easley, Jeremy Mercuri

## Abstract

Intervertebral disc (IVD) degeneration (IVDD) leads to structural and functional changes. Biomaterials for restoring IVD function and promoting regeneration are currently being investigated; however, such approaches require validation using animal models that recapitulate clinical, biochemical, and biomechanical hallmarks of the human pathology. Herein, we comprehensively characterized a sheep model of chondroitinase-ABC (C-ABC) induced IVDD. Briefly, C-ABC (1U) was injected into the L_1/2_, L_2/3_, and L_3/4_ IVDs. Degeneration was assessed via longitudinal magnetic resonance (MR) and radiographic imaging. Additionally, kinematic, biochemical, and histological analyses were performed on explanted functional spinal units (FSUs). At 17-weeks, C-ABC treated IVDs demonstrated significant reductions in MR index (p=0.030) and disc height (p=0.009) compared to pre-operative values. Additionally, C-ABC treated IVDs exhibited significantly increased creep displacement (p=0.004) and axial range of motion (p=0.007) concomitant with significant decreases in tensile (p=0.034) and torsional (p=0.021) stiffnesses and long-term viscoelastic properties (p=0.016). C-ABC treated IVDs also exhibited a significant decrease in NP glycosaminoglycan: hydroxyproline ratio (p=0.002) and changes in microarchitecture, particularly in the NP and endplates, compared to uninjured IVDs. Taken together, this study demonstrated that intradiscal injection of C-ABC induces significant degeneration in sheep lumbar IVDs and its potential for use in evaluating biomaterials for IVD repair.

**Statement of Significance:** Selecting the appropriate model for assessing biomaterials to repair and/or support regeneration of the intervertebral disc (IVD) has been controversial, leading to the use of many methods of simulating IVD degeneration (IVDD) in multiple species. Many of these models lack thorough characterization of their fidelity to human IVDD, which could hinder the translation of novel biomaterials and therapies due to unknown confounding factors. Herein, further investigation of one such model was performed using the matrix-degrading enzyme chondroitinase-ABC to induce degeneration in sheep lumbar IVDs. Degenerative changes were quantified using outcome measures relevant to human IVDD, and this dosage and method induces an aggressive degeneration environment that could be used to assess biomaterials that mimic the structure and function of the entire composite IVD. These findings may aid investigators in their selection of an appropriate animal model for preclinical testing of biomaterials and other therapeutics.

## 1. Introduction

The intervertebral disc (IVD) is a fibro-cartilaginous structure adjoining the vertebral bodies of the spine. The primary role of the IVD is to support and transmit axial loads while allowing for spinal mobility and stability during activities of daily living. Each IVD comprises three morphologically distinct regions. The centralized core of the IVD is known as the nucleus pulposus (NP); a hydrophilic, extracellular matrix (ECM) composed of aggrecan and collagen type II. The NP is circumferentially sequestered by the annulus fibrosus (AF); a ring-like structure consisting of 15-25 concentric layers of type I collagen whose fiber preferred direction is oriented at ± 28-43o to the transverse axis of the spine in alternating layers.^(1)^ This results in a fiber-reinforced composite structure with an ‘angle-ply’ microarchitecture. The third region of the IVD is known as the cartilaginous endplate (CEP), which is a thin layer of hyaline cartilage located between the inferior and superior surfaces of the IVD and adjacent vertebral bodies. The CEP serves as a mechanical barrier and allows for nutrient transport between the IVD and adjacent vertebral bodies.^(2)^

Approximately 1.5 to 4 million adults in the U.S. have IVD-related low back pain (LBP), leading to lost wages and reduced productivity exceeding $100 billion in the U.S. and $12 billion in the U.K. annually.^(3),(4)^ A common diagnosis for patients experiencing LBP is intervertebral disc degeneration (IVDD),^(5)^ which presents with several clinical characteristics. X-ray and magnetic resonance (MR) imaging often demonstrate significant reductions in overall IVD height, NP desiccation, intra-vertebral herniations (i.e. Schmorl’s nodes), changes in the vertebral body end-plates (i.e. Modic changes), and osteophyte formation.^(6)^ These changes are associated with IVDD, an aberrant, cell-mediated process originating in the NP due in part to genetic predisposition, mechanical overload, and limited nutrient supply.^(7)^ Once initiated, IVDD is marked by elevated concentrations of pro-inflammatory mediators that promote the production of ECM-degrading enzymes by resident cells.^(8),(9)^ In turn, ECM turnover favors catabolism over anabolism, eventuating the desiccation and breakdown of the NP tissue; specifically, its proteoglycan constituents. Degraded NP ECM promotes inflammation, as evidenced by increased concentrations of interleukin-1 beta (IL-1β) and tumor necrosis factor-alpha (TNF-α).^(10)^ This positive-feedback loop that couples inflammation and ECM degradation may amplify small perturbations in IVD physiology, ultimately leading to altered IVD microarchitecture and ECM mechanical properties.^(11)^ This begets altered spinal kinematics,^(12)^, and non-physiologic loading of the AF that can result in lamellar disorganization and damage leading to herniation and pain.^(13)^

The symptoms of IVDD are currently addressed by surgical discectomy to remove IVD tissue followed by joint fusion or total disc replacement. These strategies are end-stage options used to treat severe IVDD and suffer from significant limitations including promoting degeneration of adjacent levels. To address these issues and restore IVD microarchitecture and function, regenerative medicine approaches are being investigated, including the use of biomaterial scaffolds and/or mesenchymal stem cells. Employing such approaches, investigators have demonstrated the ability to promote IVD tissue regeneration *in vitro*,^(14)–(16)^ as well as the attenuation of degeneration in small animal models.^(17)^ However, biomaterial implantation within the IVD is challenging in small animal models, and clinical translation of such strategies requires evaluating their efficacy in large animal models that recapitulate the salient biochemical, mechanical, and clinical features of mild to moderate IVDD prior to human clinical trials.

Selecting the appropriate animal model to investigate the pathogenesis of IVDD and to evaluate potential biomaterials and biologics to promote IVD repair and regeneration has been controversial.^(17),(18)^ Reitmaier *et al.* reviewed literature describing studies employing large animal models used in IVD research and identified several shortcomings within the field, including minimal or no justification of animal model selection.^(17)^ Additionally, the authors suggested that logistical factors, such as behavioral differences, animal cost, availability, or researcher preference, may influence model selection in the absence of empirical differences. The authors concluded that current limitations of data concerning large animal models minimized the impact of currently published literature, warranting the further comprehensive characterization of *in vivo* models to determine their suitability for IVD research. Of those reviewed, caprine (goat) and ovine (sheep) models were found to be the most common quadrupeds used to study IVD pathology and repair.

In goats, Hoogendoorn and colleagues extensively characterized the ability of intradiscal injection of chondroitinase-ABC (C-ABC) to initiate and consistently produce progressive, degenerative changes in lumbar IVDs as evidenced by clinical imaging, biochemical, and histological analyses.^(19)–(21)^ These results were re-affirmed in a similar goat study by Gullbrand *et al.* who administered 1U C-ABC resulting in moderate IVDD by 12 weeks.^(22)^ In contrast, equally comprehensive studies have not been performed in sheep, despite significant similarities to human lumbar IVDs with respect to geometry,^(23)^ range of motion,^(18)^ ECM composition,^(24)^ age-related changes,^(25),(26)^ notochordal cell absence,^(27)^ and intradiscal pressures.^(28)^ To date, only two studies, to the authors’ knowledge, have utilized intradiscal C-ABC injection to induce IVDD in sheep.^(29),(30)^ Sasaki *et al.* performed intradiscal injections of 1, 5, or 50 U C-ABC per IVD, which resulted in significant reductions in intravital lumbar intradiscal pressures and heights over 4-weeks across all concentrations.^(29)^ Ghosh *et al.* utilized 1U C-ABC to induce degeneration over a period of 12-weeks before administering progenitor cells to evaluate their efficacy to regenerate the IVD.^(30)^ However, this study was not specifically designed to characterize the degenerative model, and only included imaging and histological outcomes. Thus, further characterization of this model using a multi-faceted approach is warranted. To address this need, the objective of the study herein was to further characterize the biochemical, mechanical, and histological changes observed in sheep lumbar IVDs following intradiscal injection of 1U C-ABC (Figure 1), and to compare the resulting degeneration to the hallmarks observed in the human pathology. The comprehensive characterization of this model will assist in determining its suitability for IVDD research and evaluating biomaterials and biological approaches for further IVD repair and regeneration.

**Figure 1.**
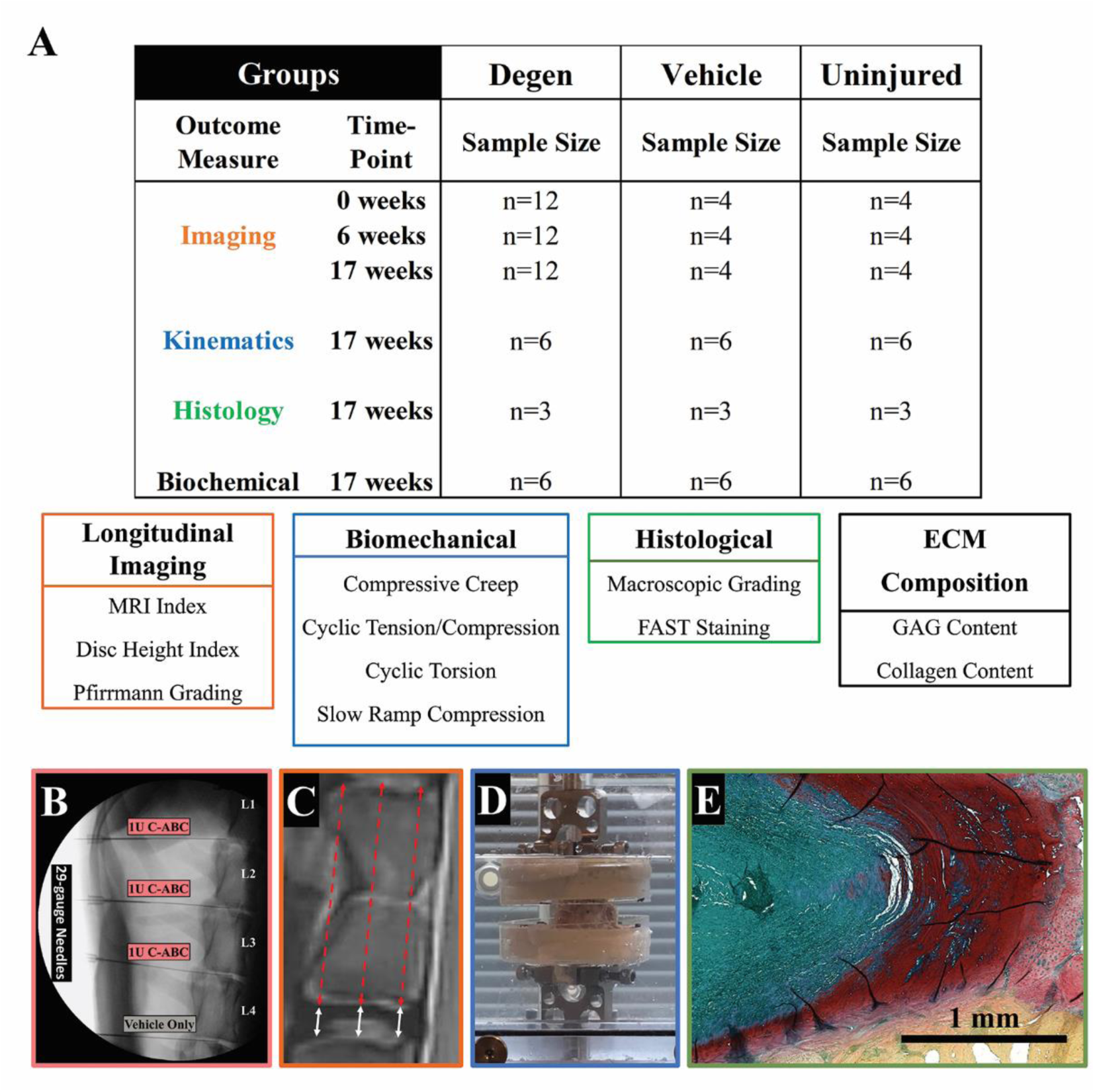
Animal study design overview and surgical approach. A) The sheep study was conducted over 17-weeks beginning with radiographic and MR imaging of healthy IVDs followed by induction of degeneration (Degen) at week 0 via intradiscal administration of 1U C-ABC. At 6- and 17-weeks, IVDs were evaluated via radiographic and MR imaging, kinematic testing, histology, and biochemical assays. B) Representative lateral fluoroscopic image illustrating image-guided intradiscal injections of 1U C-ABC in via a 29-gauge needle. C) Representative T1-weighted MR image of a sheep vertebral body and adjacent IVD demonstrating IVD (white solid arrows) and vertebral body height (red dotted arrows) measurements used for calculating DHI. D) Representative image of potted FSU undergoing kinematic testing in a saline bath. E) Representative composite histological image of an uninjured IVD stained with FAST: green /blue = NP, red = AF, and yellow = vertebral bone.

## 2. Materials & Methods

### 2.1 Institutional Review Board (IRB) Approval

The study was approved by the Institutional Animal Care and Use Committee of Colorado State University (IACUC Protocol: 16-6891A). IVDs from five lumbar levels of nine (9) skeletally mature female sheep (*Ovis Aries*, Rambouillet ewes; 62-90 kg, 3 years-of-age, K&S Livestock, Fort Collins, CO) were utilized in this study. Animals were group-housed in an indoor/outdoor pen with access to a three-sided shelter and evaluated daily by a board-certified veterinarian (J.E.) for signs of pain, behavior changes, and/or gait abnormalities for the duration of the study.

### 2.2 Surgical Procedure and Intradiscal Administration of C-ABC

All surgeries were performed by a board-certified veterinarian (J.E.) with extensive large animal spine surgery experience. Peri-operatively, a transdermal fentanyl patch (150 mcg/hr/sheep) was applied to each animal for five days starting one day before surgery. Twenty-four hours before surgery, five doses of procaine penicillin G (3 million units) and phenylbutazone (1 gram) was administered to each sheep. The animals were induced with ketamine (2mg/kg IV) and midazolam (0.2mg/kg IV), and then, intubated and maintained on 1.5-3% isoflurane in 100% oxygen throughout the surgical procedure. Under anesthesia and using standard surgical preparation, IVDD was induced via percutaneous intradiscal injection (29-gauge needle) of 1U of C-ABC (Amsbio, Cambridge, MA) in 200 µL of vehicle (sterile 0.1% bovine serum albumin in 1x PBS - Fisher Scientific, Hampton, NH) into the L_1/2_, L_2/3_, L_3/4_ IVDs (hereafter referred to as “Degen” IVDs). Before injection, IVD location and accurate needle placement were confirmed using lateral and anterior-posterior fluoroscopy. The L_4/5_ and L_5/6_ IVDs served as both Vehicle and Uninjured controls, respectively. Post-operatively, sheep were monitored until ambulatory, and then returned to standard housing conditions. Seventeen weeks following C-ABC injection, animals were euthanized by intravenous barbiturate overdose (pentobarbitone sodium, 88mg/kg) in compliance with the 2013 American Veterinary Medical Association guidelines. Immediately following euthanasia, lumbar spines were harvested *en bloc*, frozen, and shipped overnight on ice to Clemson University for further analysis. Animal IVD sample allocation are summarized in Figure 1.

### 2.3 Magnetic Resonance Imaging of IVDs

Sagittal MR imaging was performed immediately prior to intradiscal injection of C-ABC (week 0 – baseline) and tracked longitudinally in four of nine sheep at 6- and 17-weeks post-injection to monitor changes in IVDs. MR image scans were obtained using a 1.5 Tesla clinical imager (GE Signa). T2-weighted, T1-weighted, and Short-T1 Inversion Recovery (STIR) MR imaging sequences were performed on the lumbar spines. Sagittal images were constructed using a T2-weighted fast spin-echo sequence using a spine array coil (time to repetition: 2782 ms; time to echo: 101 ms; voxel size: 0.78 mm x 0.78 mm x 3.0 mm, with a 0 gap), a T1-weighted fast spin-echo sequence using a spine array coil (time to repetition: 616 ms; time to echo: 18 ms; voxel size: 0.78 mm x 0.78 mm x 3.0 mm, with a 0 gap), and a STIR sequence using a spine array coil (time to repetition: 3500 ms; time to echo: 40 ms; voxel size: 1.56 mm x 1.56 mm x 3.0 mm, with a 0 gap and inversion time of 150).

Semi-quantitative MR image analysis was performed as described by Hoogendoorn *et al*.^(21)^ MR imaging index was calculated from T2-weighted images as the product of the cross-sectional area and mean signal intensity of the encircled NP using IMPAX 6.6.1.4024 (AGFA HealthCare N.V., Mortsel, Belgium) software. Consistent mid-sagittal IVD imaging was confirmed by ensuring the full cross-sectional of the spinal cord was in view. MR imaging index is expressed as a percentage of week 0 (pre-C-ABC injection) values for normalization. Of note, normalization of signal intensity was not performed by comparing to the spinal cord, as described by others, because it is a mobile structure that can present a variable Gibbs and pulsation artifact; thus, normalization to the respective MR imaging index at week 0 was deemed most appropriate. Two researchers (R.B. and J.W.) independently quantified the initial MR imaging index for each IVD. Subsequently, a board-certified neuroradiologist (L.M.) verified and authenticated this quantitation and performed a qualitative analysis of MR images using the classification scale described by Pfirrmann *et al*.^(31)^

### 2.4 Radiographs of IVDs

IVD height index (DHI) was measured from lateral radiographs (X-Rays) and tracked longitudinally in the same four sheep at 0-, 6-, and 17-weeks post-injection to monitor changes in IVDs, in accordance with Hoogendoorn *et al.* with minor modifications.^(21)^ IVD and vertebral body heights were each quantified using three height measurements at ventral-dorsal quartiles using ImageJ (NIH, Bethesda, MD). DHI was calculated as each IVD’s mean disc height divided by the mean adjacent vertebrae height, with changes expressed as a percent of week 0 DHI. Measurements were performed separately by two researchers (R.B. and J.W.) to evaluate the interobserver reliability and were blindly authenticated by a board-certified neuroradiologist (L.M.).

### 2.5 Functional Spinal Unit Axial and Torsional Kinematics

FSU’s with intact posterior elements (n=6/group) were isolated from each spine and potted in urethane resin (Goldenwest Manufacturing, Grass Valley, CA) for testing as described previously by our group, with minor modification.^(32)^ Briefly, samples first were thawed in a test chamber filled with 1xPBS/protease inhibitor at 25°C. Once thawed, samples underwent creep loading on a Bose ElectroForce (model: 3220, TA Instruments, New Castle, DE) equipped with a 100-lb. load cell. Samples were compressed to a mean amplitude level of 0.125 MPa and then underwent a 1-hr. creep period at 0.50 MPa in compression, and then returned to a mean amplitude level of 0.125 MPa in compression before removal. Samples were then immediately transferred to a servohydraulic test frame (model: 8874, Instron, Norwood, MA) fitted with a 20kN load cell, and a mean amplitude load of 0.125 MPa in compression was applied immediately. FSU’s were subjected to 35 cycles of axial compression (0.50 MPa) and tension (0.25 MPa) at 0.1 Hz. Compression was then maintained at 0.50 MPa as samples underwent 35 torsion cycles to ± 3°. Finally, samples underwent a slow-rate compressive ramp (1 N/s) from 0.125 MPa to 0.50 MPa. A non-linear constitutive model was fit to the creep data using GraphPad Prism 7 software (La Jolla, CA) to yield elastic (Ψ) and viscous (*η*) damping coefficients for the short-term (*η*_1_ and Ψ_1_) and long-term (*η*_2_ and Ψ_2_), as described previously.^(33)^ Tensile and compressive stiffness was determined using a linear fit of the loading force-displacement curve from 60-100% of the 35^th^ cycle. Torsional stiffness was calculated from a linear fit of the loading torque-rotation curve of the 35th cycle. Torque range and axial range of motion (RoM) was calculated as the peak-to-peak torque and displacement, respectively. The constant-rate slow-ramp compression stiffness was determined using a linear fit of the
 slow-ramp load-displacement response.

### 2.6 IVD Glycosaminoglycan and Hydroxyproline (GAG:HyPro) ECM Composition

AF and NP tissues were isolated from IVDs and immediately frozen at −80°C and lyophilized to determine dry weight. Lyophilized samples were then digested in papain solution (100mM Na2PO4, 5mM N-Acetyl-L-Cysteine, 5mM EDTA, 3.875 U/mL Papain (Sigma-Aldrich)) at 65°C overnight. Digestate was then assayed for sulfated-GAG content using 1,9-Dimethyl-Methylene Blue assay with a chondroitin sulfate standard (2.5-25 µg/mL; Sigma-Aldrich).^(34)^ To quantify collagen content, digestates were hydrolyzed for 1 hour at 121°C and 15 psi in 6N HCl (Sigma-Aldrich). Hydrolyzed digestates then underwent hydroxyproline assay as described by Cissell *et al*,^(35)^ in conjunction with a hydroxyproline standard curve (5-35µg/ml; Sigma-Aldrich). Collagen content was inferred from hydroxyproline quantity using hydroxyproline: collagen percentage of 13.5%.^(36)^ Sulfated-GAG and collagen content were respectively normalized to dry weight and were additionally expressed as the ratio of sulfated-GAG to hydroxyproline (GAG:HyPro).

### 2.7 Macroscopic Evaluations and Histology of IVDs

Lumbar spines were evaluated for macroscopic and histological changes. FSUs were disarticulated from each spine via bandsaw and fixed for 7 days in 10% neutral-buffered formalin, followed by decalcification in 12% formic acid. Samples were prepared into 3-mm sagittal slices and images were captured for macroscopic evaluation by 3 blinded observers (R.B., J.W., and J.M.) using the Thompson grading scale (Supplemental 1–3).^(37)^ These mid-sagittal slices were paraffin-embedded, and then, 7µm sections were obtained. Staining was performed *en masse* and in accordance with methods from Leung *et al*.^(38)^ Sections were rehydrated via ethanol gradations and stained with 3% Alcian blue (pH 1.0) for 8 min., 0.1% safranin-O for 6 min., 0.25% tartrazine for 10 sec, 0.001% fast green for 4 min. Micrographs were acquired using an Axio Vert.A1 microscope with AxioVision SE64 Rel. 4.9.1 software (Zeiss, San Diego, CA). Composite images were stitched together using FIJI analysis software.^(39),(40)^ Composite IVD micrographs were scored using the IVD degeneration scale described by Walter *et al.* with minor modification (Supplemental 4–6).^(41)^ Endplate Integrity, AF Morphology, AF/NP Demarcation, NP Matrix Homogeneity, and NP Matrix Stain Intensity were each scored from 0 to 2 by two blinded observers (RAB, JJM) and an unblinded observer (JDW). The mean score for each category was summed to produce a semi-quantitative aggregate score.

### 2.8 Statistical Analysis

Results are represented as mean ± standard error of the mean (SEM) and significance was defined as (p<0.050). Statistical analysis was performed using GraphPad Prism 7 software. Comparisons were performed using a one-way ANOVA with Holm-Sidak method for multiple comparisons (MR imaging Index, DHI, ECM Composition) or Dunnett’s post-hoc (FSU kinematics) analysis. Due to high observed variability, outlier analysis was performed on ECM composition data using the ROUT method (Q = 1%).^(42)^ Pfirrmann scoring was evaluated via a Kruskal-Wallis test with Dunn’s multiple comparisons. Inter-observer reliability of Thompson grades and Rutges scores was evaluated using IBM Statistical Package for Social Sciences (SPSS 24.0; IBM, Armonk, NY). The Intra-class Correlation Coefficient (ICC) was calculated using a two-way random model for absolute agreement as described by McGraw and Wong.^(43)^ An ICC of 0.4-0.75 indicates good agreement, while >0.75 is considered excellent.^(44)^ Mean Thompson and histological scores were compared between groups via a one-tailed Mann Whitney test.

## 3. Results

Eight of the nine animals tolerated the intradiscal C-ABC injection procedure, recovered from anesthesia, and began weight-bearing and eating within one hour postoperatively. One animal had difficulty standing after surgery, however, the animal progressively improved and exhibited good mobility and no evidence of weakness.

### 3.2 Intradiscal C-ABC Injection Results in Significantly Reduced MR Imaging Index

To determine the effect of C-ABC injection on IVD hydration, MR imaging was performed and accompanied by MR index calculations and Pfirrmann scoring of T2-weighted images. Mean MR index values of Degen IVDs demonstrated significant decreases by 17 weeks, reaching 75.52 ± 22.79%; (p=0.075) and 67.65 ± 16.63%; (p=0.030) of their week 0 values at 6- and 17-weeks post-injection, respectively (Figure 2). Moreover, mean MR index values for Degen IVDs were significantly lower at 17 weeks compared to Uninjured IVDs (104.28 ± 10.45%; p=0.033) at the same timepoint. No significant differences in MR imaging index values were found comparing Uninjured and Vehicle control IVDs. Qualitatively, T2-weighted MR images illustrated darkening in the NP-region of Degen IVDs (Figure 2). Of note, darkening in vertebral bodies adjacent to Degen IVDs was also observed on MR images (Figure 2), which conversely, were not evident in Uninjured and Vehicle IVDs. Degen IVDs also demonstrated an increase in Pfirrmann grade indicating worsening degeneration at week 17 (1.25 ± 0.25); however, this was not significantly different compared to Uninjured (1 ± 0) and Vehicle controls (1 ± 0) nor the IVDs respective week 0 values (1 ± 0).

**Figure 2.**
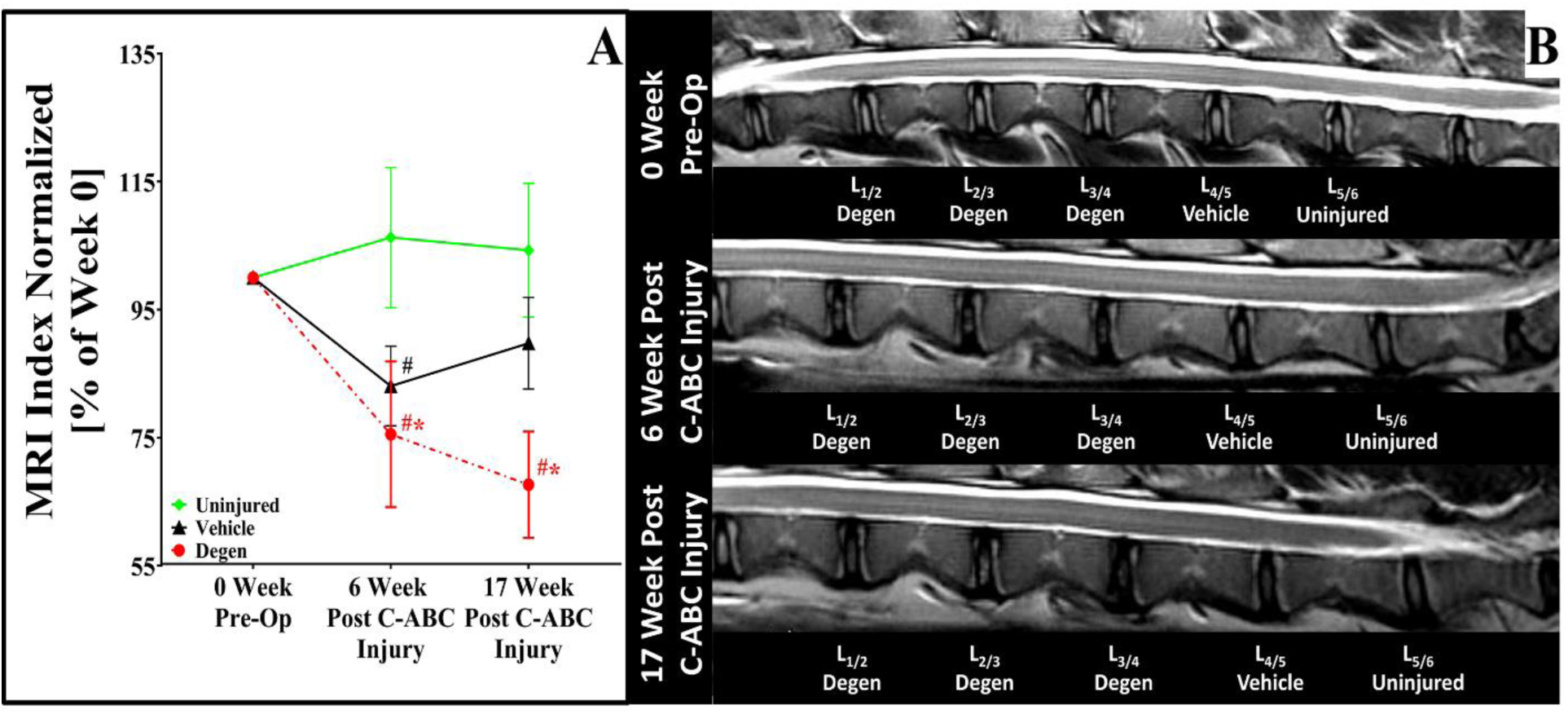
Longitudinal MR image tracking of sheep lumbar IVDs. A) Graph depicting the normalized MR image index. B) Representative longitudinal T2-weighted MR images of a single sheep throughout the 17-week study. (*) indicates a significant difference (p<0.05) between groups within the same time-point. (#) indicates a significant difference (p<0.05) within the study group compared to the pre-op (week 0) time-point.

### 3.2 Intradiscal C-ABC Injection Results in Significant Reductions in IVD Height

To determine the effect of C-ABC injection on IVD height, sagittal x-rays were obtained and disc height index (DHI) was calculated. DHI values of Degen IVDs significantly decreased at weeks 6 and 17, reaching 73.34 ± 5.74% (p=0.031) and 70.35 ± 4.16% (p=0.009) of their week 0 values, respectively (Figure 3). These values were also significantly lower compared to Uninjured IVD values at week 6 (96.17 ± 4.04%; p<0.001) and week 17 (90.40 ± 1.34%; p<0.001), respectively. Uninjured IVDs illustrated a significant decrease in DHI at week 17 (p=0.009) compared to week 0 values. Furthermore, Vehicle IVDs illustrated a significant decrease by weeks 6 and 17, reaching 83.84 ± 3.37% (p=0.028) and 81.01 ± 0.99% (p<0.001) of their week 0 values, respectively (Figure 3). Additionally, these values were significantly lower compared to Uninjured IVD values at week 6 (96.17±4.04%; p=0.013).

**Figure 3.**
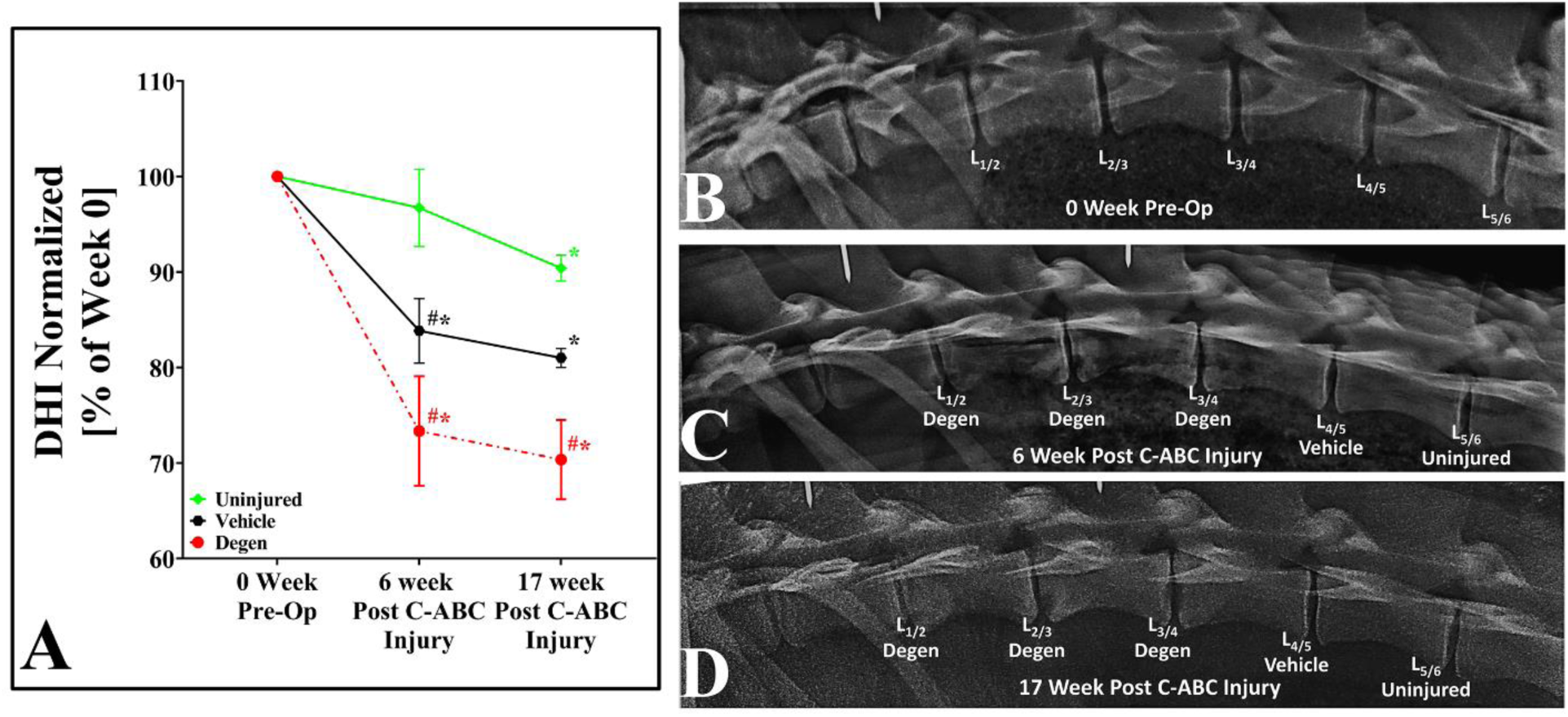
Longitudinal radiograph image tracking of sheep lumbar IVDs. A) Graph depicting disc height index (DHI). B) Representative longitudinal radiographic images of a single sheep throughout the 17-week study. (*) indicates a significant difference (p<0.05) between groups within the same time-point. (#) indicates a significant difference (p<0.05) within the study group compared to the pre-op (week 0) time-point.

### 3.3 Intradiscal C-ABC Injection Results in Significant Alterations in Functional Spinal Unit Kinematics

To evaluate the effect of C-ABC injection on FSU function, axial and torsional kinematic testing was performed. Kinematic creep loading (Figure 4) of Degen IVD FSUs demonstrated a significantly increased creep displacement (0.49 ± 0.04 mm; p=0.004) compared to Uninjured IVD FSUs (0.33 ± 0.03 mm, respectively) (Figure 4D). Additionally, long-term elastic (Ψ_2_: 537.40 ± 36.05 N/mm; p=0.016) and viscous (η_2_: 1.33 ± 0.18 x10^6^Ns/mm; p=0.016) damping coefficients of Degen IVD FSUs were significantly lower compared to Uninjured IVD FSUs (Ψ_2_: 766.75 ± 69.98 N and η: 2.07 ± 0.19×10^6^ Ns, respectively) (Figure 4E&F). No significant differences were noted in creep kinematic parameters between Uninjured and Vehicle IVD FSUs.

**Figure 4.**
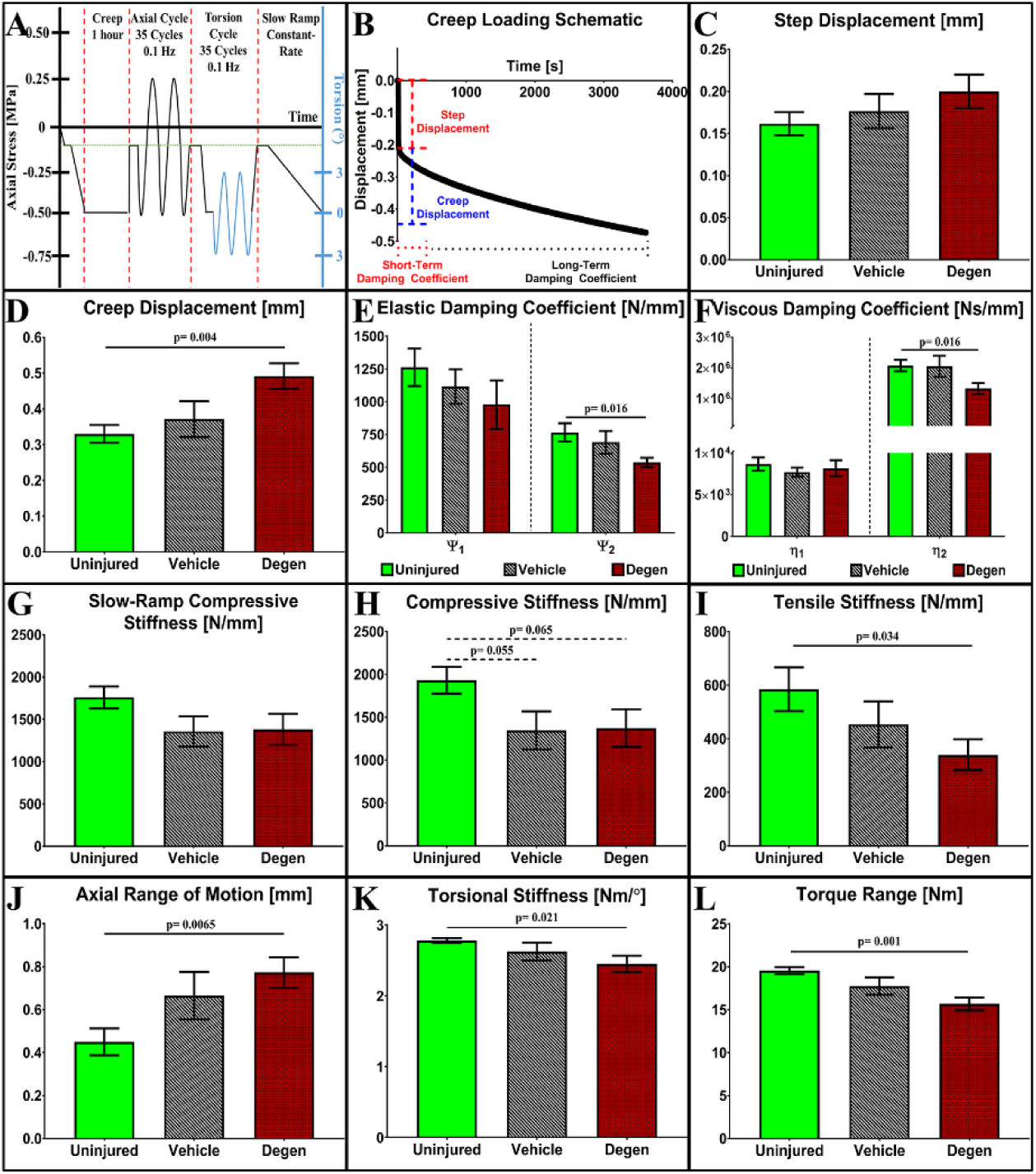
Kinematic testing of functional spinal units. A) Loading scheme for FSU testing depicting creep, axial cyclic tension-compression, axial torsion, and slow constant-rate ramp compression testing. B) Representative graph depicting the creep response of a sheep FSU and associated creep parameter measures. Graphs depicting C) step and D) creep displacement, E) elastic and F) viscous damping coefficients, G) slow ramp compressive stiffness, H) cyclic compressive and I) tensile stiffnesses, J) axial range of motion, K) torsional stiffness, and L) torque range values of Uninjured, Vehicle and Degen sheep FSUs, respectively. Solid lines connecting groups indicates a significant difference (p≤0.050).

Axial cyclic kinematic testing of Degen IVD FSUs demonstrated a significant decrease in tensile stiffness (339.60 ± 58.46 N/mm; p=0.034) and a significant increase in the axial range of motion (0.77 ± 0.07 mm; p=0.007) compared to Uninjured IVD FSUs (584.71 ± 81.70 N/mm and 0.45 ± 0.06 mm, respectively) (Figure 4I&J). Compressive stiffness values for Degen IVD FSUs (1372.60 ± 221.50 N/mm; p=0.065) and Vehicle (1347.10 ± 331.36 N/mm; p=0.055) IVDs were lower than Uninjured IVD FSUs (1933.70 ± 155.52 N/mm) but did not reach statistical significance (Figure 4H). Torsional rotation of Degen IVD FSUs demonstrated a significant decrease in torque range (15.68 ± 0.75 Nm; p= 0.001) and torsional stiffness (2.45 ± 0.12 Nm/°, p=0.021) compared to Uninjured IVD FSUs (19.57 ± 0.41 Nm and 2.78 ± 0.03 Nm/°, respectively) (Figure 4K&L). No significant changes were observed between any torsional kinematic parameters of the Uninjured and Vehicle IVD FSUs.

### 3.4 Intradiscal C-ABC Injection Results in a Decrease in NP and AF GAG:HyPro ratio

To determine the effect of C-ABC injection on NP and AF ECM composition, changes in glycosaminoglycan and hydroxyproline (GAG:HyPro) content were evaluated. A two-way ANOVA indicated animal variability (p=0.027), tissue specificity (p<0.001), and animal-tissue interactions (p=0.022) contributed significantly to the observed variability in the biochemical composition. A single outlier was found and removed from the collagen content data (Degen-NP), and non-significant trends in the GAG and collagen content from AF and NP tissues showed marginal increases in collagen with concurrent decreases in GAG content (Figure 5). The GAG:HyPro ratio in the NP region of Degen IVDs was significantly lower (6.75 ± 1.77) compared to Uninjured IVDs (16.52 ± 1.89; p=0.002) and Vehicle IVDs (14.16 ± 1.97; p=0.014); moreover, the ratio of GAG:HyPro was decreased in the AF region of Degen IVDs compared to Uninjured IVDs.

**Figure 5.**
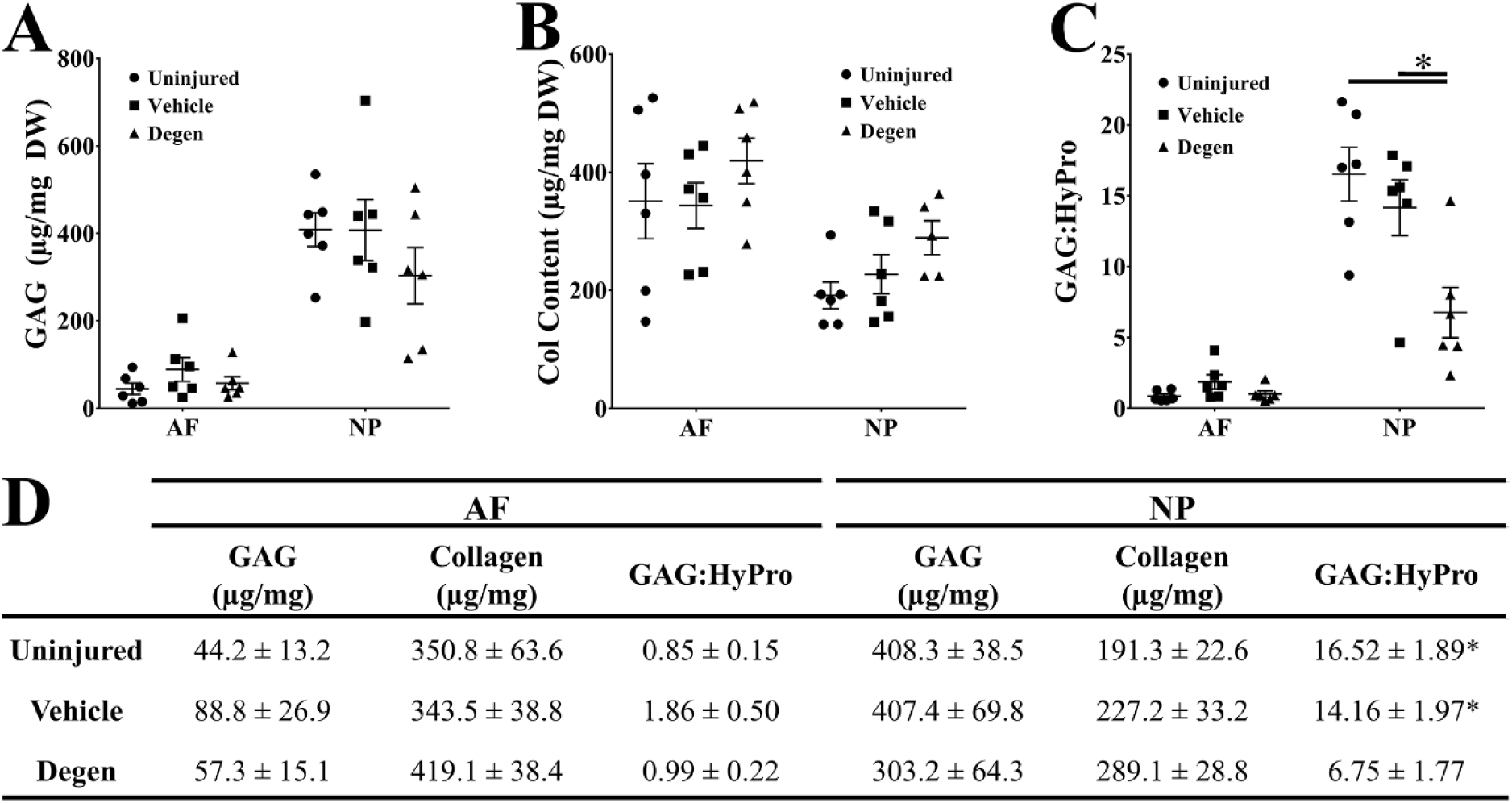
Extracellular matrix composition of IVD tissues. Graphs illustrating A) GAG and B) collagen content of AF and NP tissues isolated from Uninjured, Vehicle, ad Degen FSUs. Graph depicting C) computed GAG: Hydroxyproline (HyPro) ratio. D) Tabulated ECM composition data expressed as mean ± SEM. (*) indicates a significant difference (p≤0.050) between groups connected by horizontal bars.

### 3.5 Intradiscal C-ABC Injection Induced Significant Changes in IVD Morphology and Microarchitecture

To determine the effect of C-ABC injection on IVD structure and microarchitecture, macroscopic images and FAST stained histological sections were assessed via Thompson and Rutges grading, respectively (Figures 6 and 7). Thompson grading of macroscopic images showed perfect agreement (ICC: 1.0; Supplemental 3) and reflected significant degenerative changes in Degen IVDs. Uninjured and Vehicle IVDs had mean Thompson grades of 1.0 ± 0.0, while Degen IVDs were significantly greater (4.0 ± 0.0) than Uninjured (p=0.014) and Vehicle (p=0.014) IVDs (Figure 6B). Initial macroscopic evaluation of Uninjured and Vehicle IVDs showed features indicative of healthy IVDs, such as turgid and white NPs, crescent-shaped AF lamellae, and continuous CEPs with uniform thickness (Supplemental 2). Degen IVDs had significant alterations to macroscopic features compared to Uninjured and Vehicle IVDs. Macroscopically, the NP regions of Degen IVDs exhibited a darkened and irregular matrix, and AF regions showed disorganization. Critically, CEP integrity and uniformity were diminished, as evidenced by multiple protrusions into the subchondral bone along the endplate.

**Figure 6.**
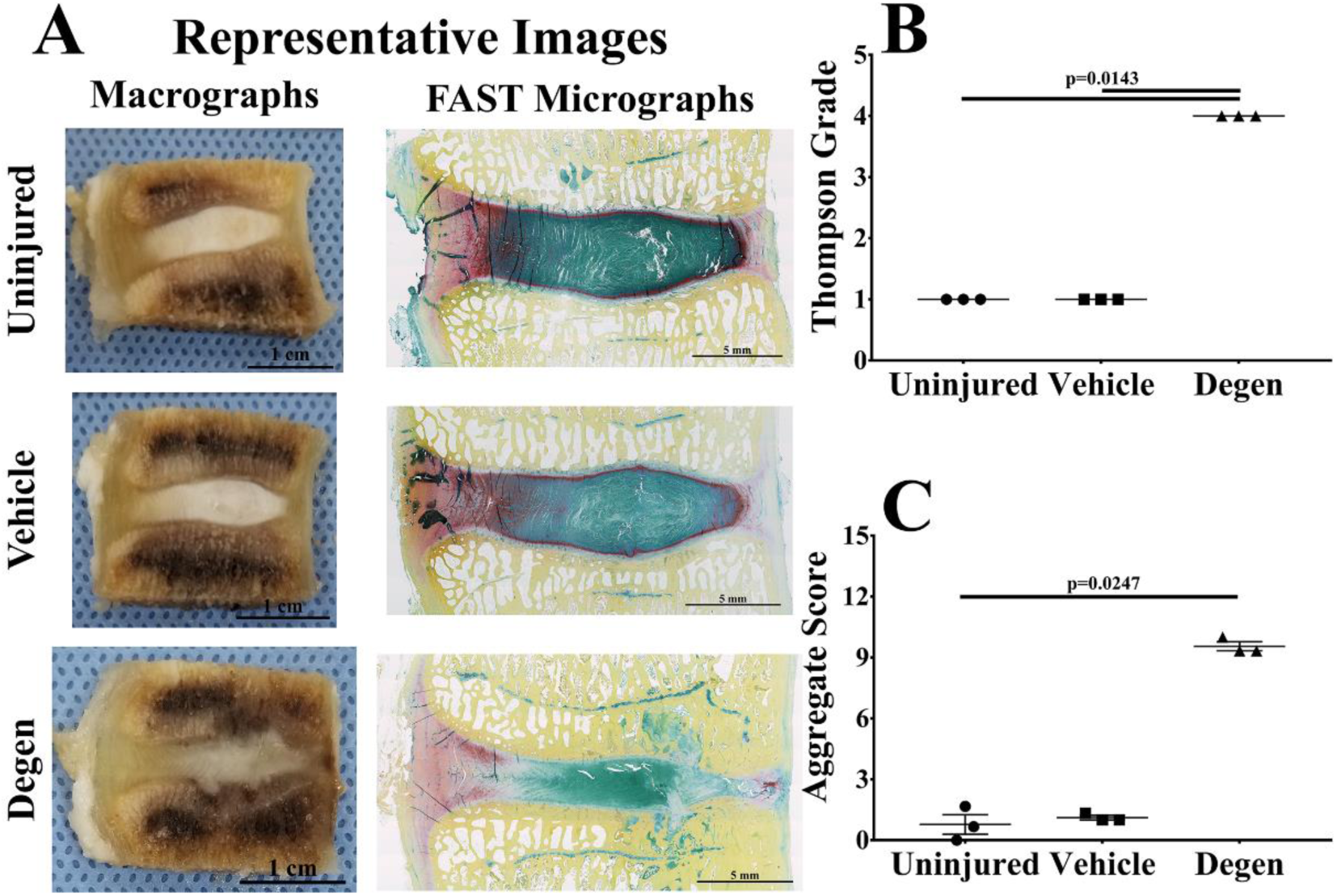
Morphological changes in IVDs. A) Representative macrographs and micrographs of Uninjured, Vehicle, and Degen IVDs from a single animal. B) Graph depicting Thompson grading of macrographs prior to histological processing. C) Graph depicting Rutges scoring applied to FAST-stained micrographs. Light Green: Outer AF; Dark Red: Inner AF; Teal: NP; Yellow: Vertebral Bone. Bars connecting groups indicate significant differences (p≤0.050).

**Figure 7.**
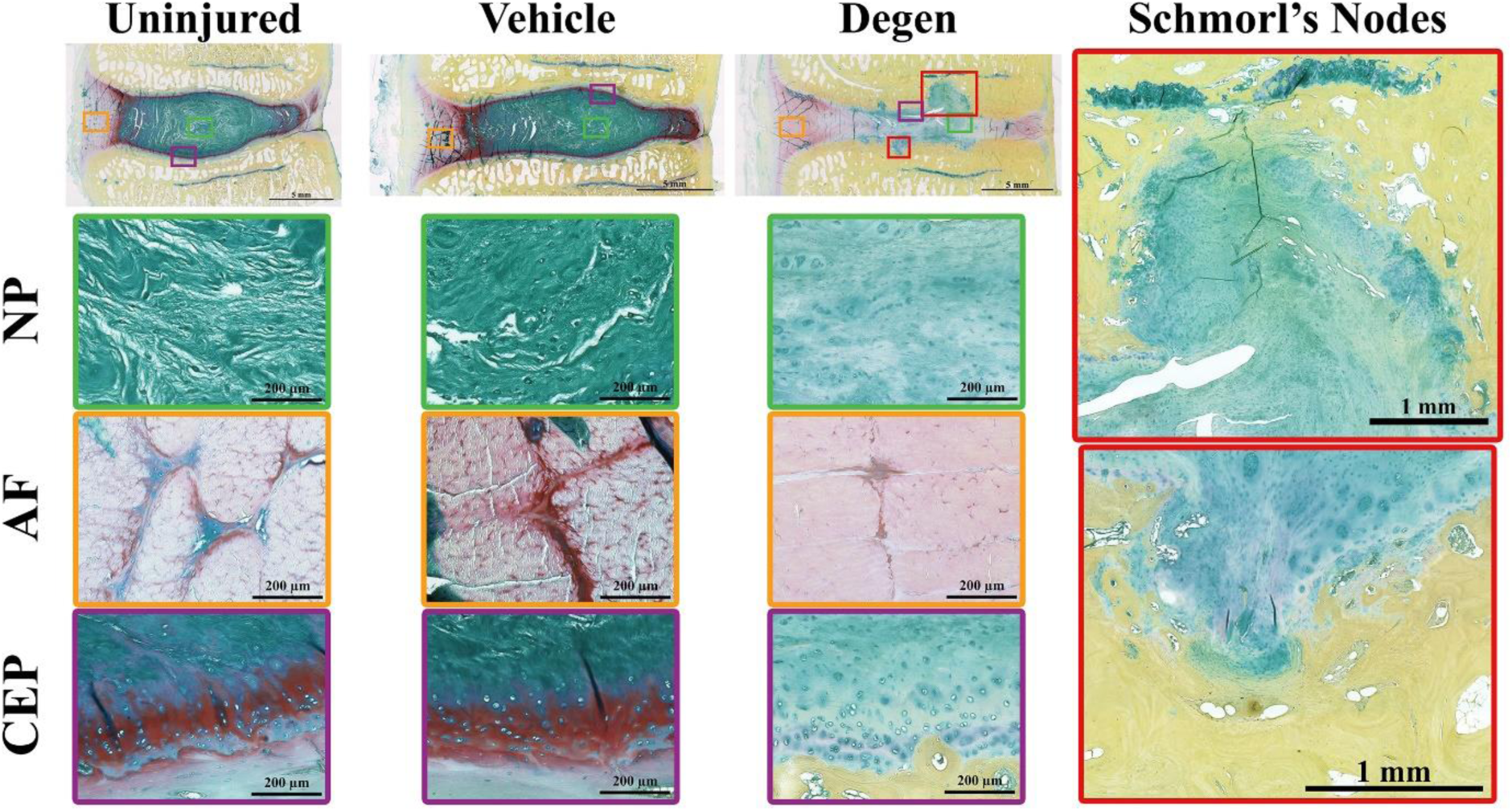
Histological features of a representative set of FAST-stained micrographs. Representative histological features from the NP, AF, and CEP regions of Uninjured, Vehicle and Degen IVDs as well as representative images depicting Schmorl’s nodes observed in Degen IVDs.

Following histologic sectioning and staining, semi-quantification of histomorphological changes using the Rutges scale showed excellent agreement (ICC: 0.94; Supplemental 6) among observers. Uninjured and Vehicle IVDs received mean aggregate scores of 0.8 ± 0.5 and 1.1 ± 0.1, respectively. (Figure 6C), In contrast, the mean aggregate score for Degen IVDs (9.6 ± 0.2) was significantly higher than Uninjured IVDs (p=0.025) and marginally higher than Vehicle IVDs (p=0.072), indicating a presence of higher degree degeneration. NP regions of Degen IVDs showed notable heterogeneity and diminished stain intensity, while AF regions demonstrated derangement of the multi-lamellar structure and inversion of inner AF layers into the NP (Figure 7 - **NP and - AF**). Furthermore, the distinction between AF and NP was markedly reduced. The CEP was strikingly altered, with an almost total absence of staining compared to Uninjured and Vehicle controls (Figure 7 - **CEP**). Additionally, CEP integrity was notably compromised, as numerous protrusions into the subchondral bone were observed. Most prominently, Schmorl’s nodes encroaching approximately 2-3 mm into the subchondral bone were observed, and bone adjacent to these nodes appeared denser than that of control IVDs. (Figure 7 - **Schmorl’s Nodes**)

## 4. Discussion

In this study, the morphological, mechanical, biochemical, and histological changes of C-ABC induced IVDD was characterized in sheep lumbar IVDs. Findings of this study indicate intradiscal injection of 1U of C-ABC produced intradiscal changes resulting in severe degenerative changes by 17 weeks. This was indicated by significant alterations in clinical imaging characteristics of IVDs and adjacent vertebral bodies, FSU axial kinematics, NP GAG:HyPro ratio, and IVD microarchitecture including derangement of the NP and AF as well as formation of CEP irregularities.

C-ABC is a bacterial derived proteolytic enzyme with high specificity for chondroitin sulfate; the primary constituent of aggrecan within the NP. Specifically, C-ABC has been previously evaluated for its therapeutic potential to degrade tissue fragments impinging on nerves in patients with herniated IVDs.^(45)^ The administration of C-ABC enzyme initiates the targeted degradation of NP tissue to provide an accelerated degeneration cascade similar to conditions observed in humans as a result of increased levels of matrix metalloproteinases (MMPs) and a disintegrin and metalloproteinase with thrombospondin motifs (ADAMTS) observed within increasing levels of IVDD; thus, intradiscal injection of C-ABC has been used as a method of initiating IVDD in large animals to serve as models for studying pathological mechanisms and preclinical assessments of potential therapeutics.^(21),(22),(29),(30)^.

Herein, clinical MR imaging of C-ABC injected sheep lumbar IVDs demonstrated significant reductions in T2-weighted MR index values and signal intensity within the NP region and adjacent vertebral bodies of C-ABC treated IVDs over 17 weeks. Clinically, T2-weighted imaging provides a hyperintense (e.g. bright) image in hydrated tissues, including the IVD, which conversely appear dark on T1-weighted images. Furthermore, the correlation between T2-weighted MR imaging parameters and IVD water content has been established previously.^(46)^ Degenerate IVDs demonstrate varying degrees of hypointensity (e.g. darkening or “black”) and reduced T2 relaxation times indicative of reductions in water content. Similar findings have been observed by others after intradiscal administration of 1U C-ABC in goat and sheep IVDs.^(22),(30)^ However, one notable difference between this study and others was the observed hypointense T2-weighted signal within vertebral bodies adjacent to Degen IVDs; which was not specifically mentioned by others.^(22),(30)^ Such changes have been observed clinically in humans and often represent a spectrum of biological alterations within vertebral bodies including fatty marrow changes, endplate sclerosis, trabecular fracture, or edema and inflammation.^(47)^ In contrast to the MR index, no significant differences were observed in the Pfirrmann scores; however, this may be due to limitations of the scoring system, which only captures NP/AF features, and therefore, overlook the severe changes observed in the vertebra and endplates. Moreover, morphological differences between ellipsoid-shaped human discs and sliver-like sheep discs further complicate the scoring process. Finally, IVD heights decreased by approximately 29% at 17 weeks in IVDs receiving C-ABC. Such findings are also observed clinically in patients with IVDD indicating the inability of the degenerate IVD to support axial loads.^(1)^ Together, this can lead to foraminal stenosis, nerve impingement, facet arthropathy, and pain. Similar to our results, reductions in IVD heights of 19% - 40% have been observed by others at approximately 12 weeks post-intradiscal injection of 1U C-ABC in sheep and goats.^(22),(29),(30)^

Accompanying desiccation and reductions in IVD height, axial and torsional mechanics of sheep FSUs were also detrimentally impacted by C-ABC injection. Significant increases in creep displacement and axial range of motion in conjunction with decreases in short and long-term elastic damping coefficients, long-term viscous damping coefficients, and compressive stiffness values were noted compared to Uninjured IVDs. This has also been observed in degenerate human IVDs that exhibit increased range of motion,^(48),(49)^ depressurization of the NP,^(50),(51)^ reduced NP modulus,^(52)^ and decreased late creep responses.^(33)^ Furthermore, diminished early elastic creep responses have been noted in nucleotomized sheep IVDs, which has been shown to be dependent upon the extent of NP tissue removal.^(53)^ Ultimately, findings herein are in alignment with others demonstrating that intradiscal administration of 1U C-ABC in goat IVDs resulted in the increased neutral zone and creep displacements concomitant with reduced compressive modulus 12 weeks post-injection.^(22)^

Detrimental changes in AF and NP tissue biochemistry were also noted in C-ABC treated IVDs. More specifically, the NP GAG:HyPro ratio of Degen IVDs were significantly lower compared to Uninjured and Vehicle IVDs. Human NP tissue also demonstrates a reduction in the GAG:HyPro ratio with increasing grades of degeneration, which is approximately 27:1 in healthy IVDs compared to 5:1 in severely degenerate IVDs.^(54)^ Data herein suggest that NP GAG content decreases while its collagen content increases relative to dry weight following C-ABC injection; thus, the IVD is becoming more fibrotic. It is likely that these ECM changes represent a breakdown of aggregating proteoglycan within the C-ABC injected IVDs, which would result in the subsequent loss of osmotic potential and increased permeability.^(51),(52),(55),(56)^ Together, these alterations have been shown to contribute to the diminished ability of the IVD to effectively support compressive loading, and therefore, leading to a reduction in IVD height and overall mechanical dysfunction, as observed herein.^(6),(11),(57)^

Semi-quantitative and qualitative histological evaluations of Degen IVDs demonstrated significant alterations in the NP, AF, and CEP disruptions as compared to Vehicle and Uninjured IVDs. This was demonstrated by decreases in proteoglycan staining intensity and disorganization of the NP, alterations in the lamellar architecture of the AF, and the formation of Schmorl’s nodes (breaches of the NP into adjacent vertebral bodies), respectively. Taken together, these histomorphological features are suggestive of severe IVDD and support the imaging, mechanical and biochemical findings herein. In comparison, aged and degenerate human IVDs have been reported to display similar histological changes including increases in mucoid degeneration, cleft formation and fibrotic changes within the NP, loss of lamellar concavity, and increased presence of radial tears and rim lesions in the AF, and in addition, thinning and loss of continuity of the CEP and sclerosis of the vertebral body bone.^(58),(59)^ Of note, histological evaluations of the sheep IVDs herein clearly illustrated the penetration of NP tissue into clefts within the CEP reminiscent of Schmorl’s nodes found in human IVDs. It is hypothesized that C-ABC degraded proteoglycan within the NP and CEP. In fact, herein CEP thickness was found to be irregular upon both macroscopic and microscopic analyses. Moreover, the formation of these Schmorl’s nodes could account for the hypointense signals observed on MR imaging in adjacent vertebral bodies of C-ABC injured IVDs. The CEP normally functions to prevent the NP from breaching the vertebral body and prohibits contact with bone marrow and the immune system. This is important as extradiscal NP tissue has been shown to be autoimmunogenic,^(60)^ and thus, the observed hypointense signal in our T2-weighted MR likely represent edema and inflammation. However, further investigation is needed to confirm. Of note, CEP breaches were not observed in vehicle IVDs, which further suggests that C-ABC induced initial CEP damage as opposed to creating excessive pressurization of the IVD due to injection volume. In the context of human lumbar IVDD, the presence of Schmorl’s nodes have been shown to be correlated with Modic changes and grade of degeneration in human IVDs.^(61),(62)^ Furthermore, results herein are also in agreement with other studies in which the loss of CEP structure and protrusions of NP material into underlying bone led to vertebral bone remodeling as observed following intradiscal administration of 1U C-ABC in goats.^(22)^

An interesting finding in this study is that in the absence of morphological changes in Uninjured and Vehicle groups, when significant reductions in DHI at 17 weeks were observed. Moreover, Vehicle IVDs demonstrated kinematic parameters comparable to Degen IVDs or intermediate values between Degen and Uninjured IVDs. This trend was also observed for the collagen content and GAG:HyPro ratio in the NP of Vehicle IVDs. A probable explanation is adjacent segment degeneration arising from multi-level disc degeneration induced in the L_1/2_, L_2/3_, and L_3/4_ IVD levels. Cheung *et al.* investigated the prevalence and risk factors associated with multi-level (≥2) disc degeneration in a cross-sectional cohort and suggested that the biomechanical deficiencies of degenerative IVDs arising from altered biochemical and structural integrity may re-direct forces in the spine to adjacent, non-degenerated disc levels.^(63)^ This compensatory redistribution of spinal loads to non-degenerative IVDs could then contribute to contiguous multi-level disc degeneration through overloading and aberrant kinematics. This mechanism could explain the mild, graded degenerative trends in Vehicle and Uninjured groups, since Degen IVDs demonstrated biomechanical deficiencies indicative of joint laxity and diminished load-bearing capacity, including increased RoM and creep displacement with a concurrent decrease in torque range, stiffness, and long-term creep parameters.

As with any study, limitations were noted. First being the selection of the respective groups within each animal used in this study. Herein, three consecutive IVDs were injected with C-ABC, and the location of IVDs receiving C-ABC injection and those serving as controls were not randomized to account for differences due to spine level across multiple sheep. However, data from each IVD was normalized to its respective week 0 (baseline) value to address this. Second, while an established T2-weighted MR imaging modality was used to evaluate ECM degradation as a function of IVD hydration, it would have been advantageous to include a T1-ρ relaxation parameter, which may correlate more strongly with changes in IVD GAG content and histology.^(64)^ Additionally, varying doses of C-ABC were not evaluated, which has been shown by others to result in different degrees of degeneration in sheep.^(22)^ However, the goal herein was to provide further in-depth characterization employing a common C-ABC dosage used previously in sheep by others. Thus, further investigations using lower C-ABC concentrations could result in milder and more moderate IVDD. Additionally, it should be noted that C-ABC is not a naturally occurring enzyme found in humans and that MMP-s and ADAMTS’s drive degeneration. However, our goal herein was to detail the degenerative changes in a more commonly employed model. Finally, with the exception of clinical imaging, we only characterized the ensuing degenerative changes at ~ 4-months post injury as compared to evaluating all outcome measures longitudinally. However, the time-point evaluated is equivalent to- or exceeds those used by others. Moreover, our results demonstrate that spontaneous improvement / regeneration of the injured IVDs does not occur during this time and thus is a relevant timeframe to assess repair strategies employing biomaterials.

## 5. Conclusions

In conclusion, intradiscal administration of 1U C-ABC per lumbar IVD resulted in many of the salient hallmarks observed in severe IVDD by 17 weeks. Thus, this model can be useful in evaluating biomaterials used to restore IVD function.

## Acknowledgments

The authors would like to acknowledge the veterinary staff at Colorado State University, Dr. Patrick Gerard for his assistance with statistical analysis, and Prisma Health Upstate for allowing us to use their IMPAX Radiology Information System. Funding for this research was provided in part by the National Institute of General Medical Sciences of the National Institutes of Health (5P20GM103444), and the Clemson University Research Foundation Technology Maturation Fund. RB was supported by the National Science Foundation Graduate Research Fellowship (Award number: 2011382).

## Author Contributions

Study design (R.B., J.W., S.G., J.M., L.M., J.E.), surgical implementation (J.E.), sample processing (R.B., J.W., A.M.), data analysis (R.B., J.W., J.M., A.M., L.M.), manuscript preparation (R.B., J.W., J.M.). All authors have read and approved the final submitted manuscript.

---

**Supplement 1.**
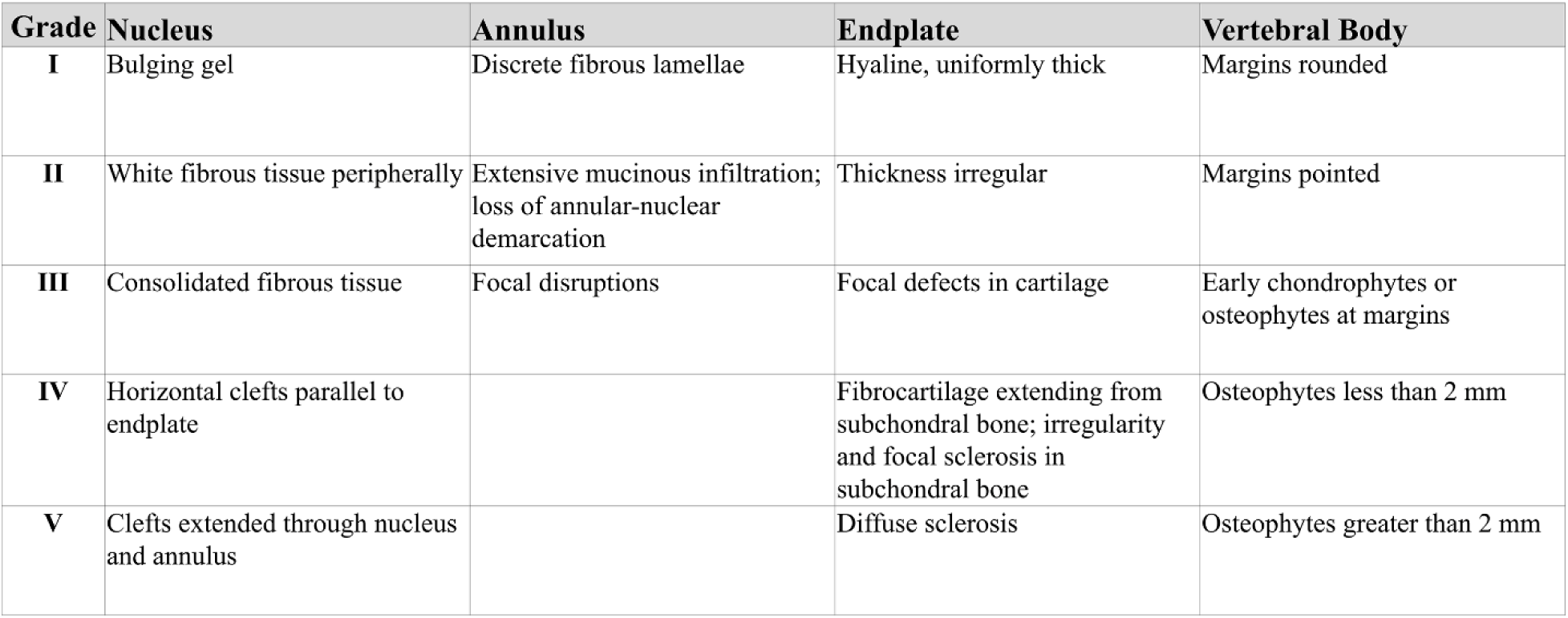
Thompson criteria used for macroscopic grading. Changes in macroscopic morphology were semi-quantitatively evaluated using the grading scale developed by Thompson *et al.* Based on the criteria above, a grade of I-V was given to each region of the motion segment for the nucleus pulposus, annulus fibrosus, cartilaginous endplate, and vertebral bodies. The average grade across all regions was then calculated and assigned to the IVD.

**Supplement 2.**
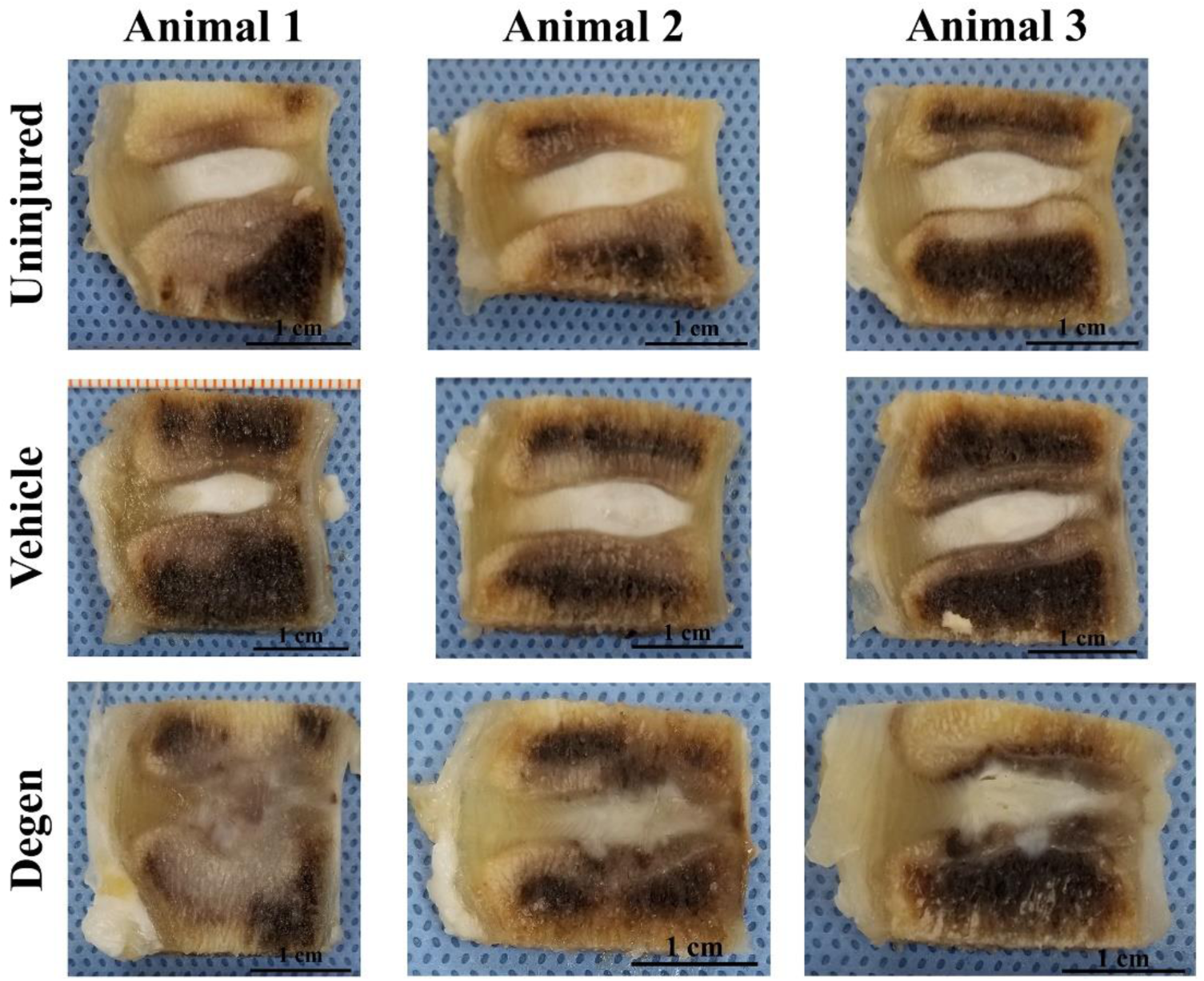
Mid-sagittal images used for Thompson grading. Uninjured, Vehicle, and Degen IVDs were isolated from three respective animals for analysis. Following formalin fixation and decalcification, mid-sagittal slices (~3mm thickness) were imaged and graded using the Thompson criteria.

**Supplement 3.**
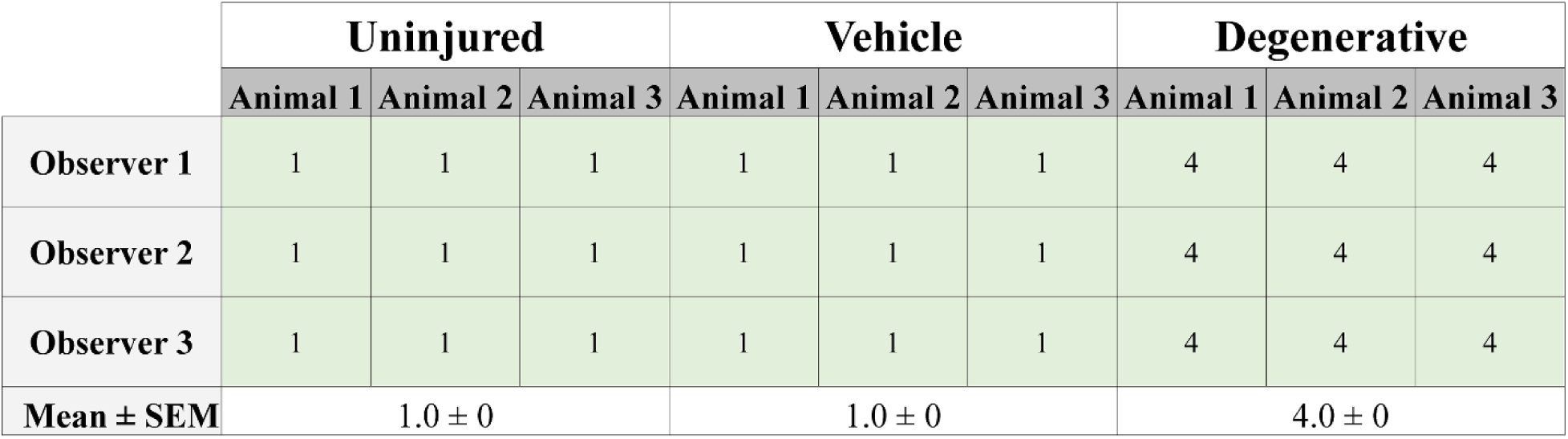
Raw Thompson grades for each animal and observer. Thompson grading for each sample was performed independently by three blinded observers to determine the inter-observer reliability, which showed perfect agreement (ICC:1.00). Grades were subsequently averaged across observers for further analysis.

**Supplement 4.**
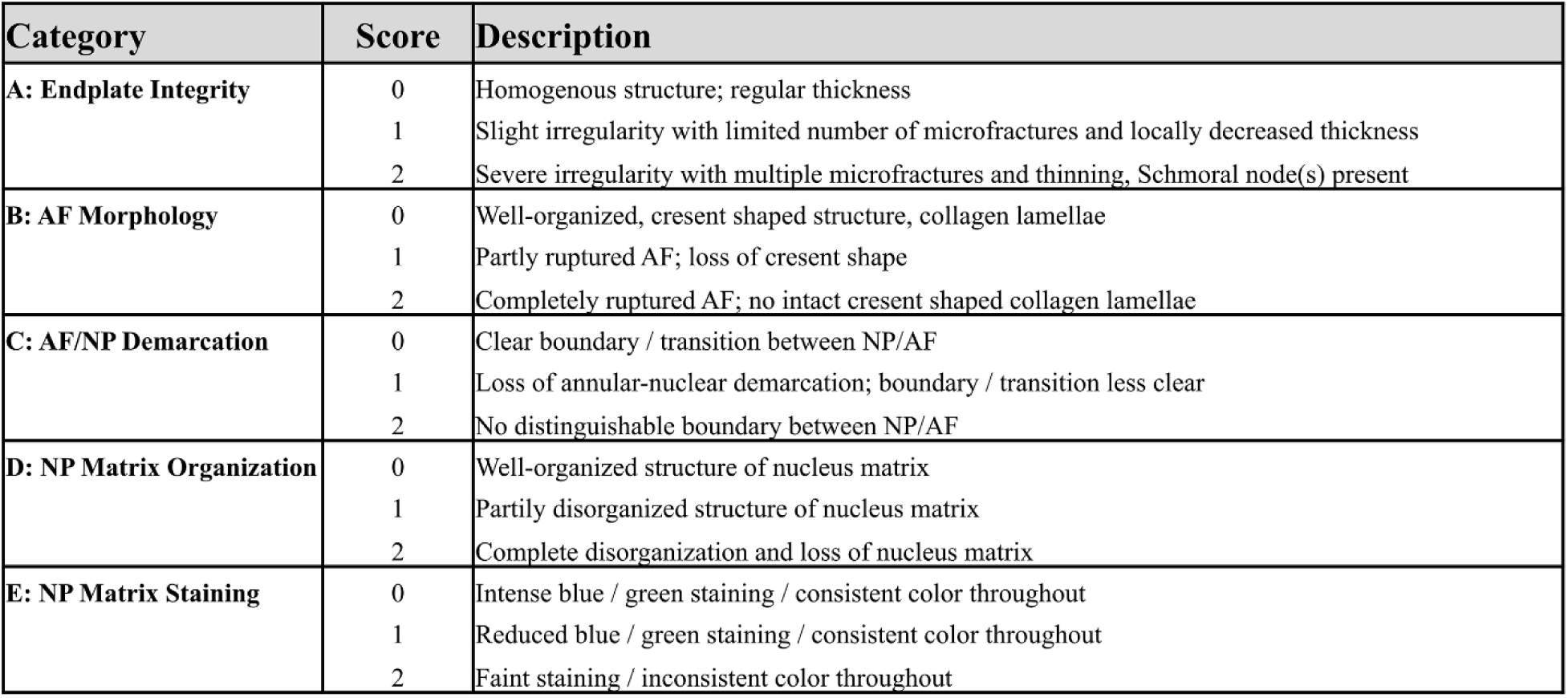
Rutges criteria used for histological scoring. Changes in morphology and microarchitecture were semi-quantitatively evaluated using the scoring criteria described by Rutges *et al.* with minor modification. Endplate integrity, AF morphology, AF/NP demarcation, NP matrix organization, and NP matrix staining were each scored 0-2 based on the above descriptors with an increasing score indicating worsening degeneration.

**Supplement 5.**
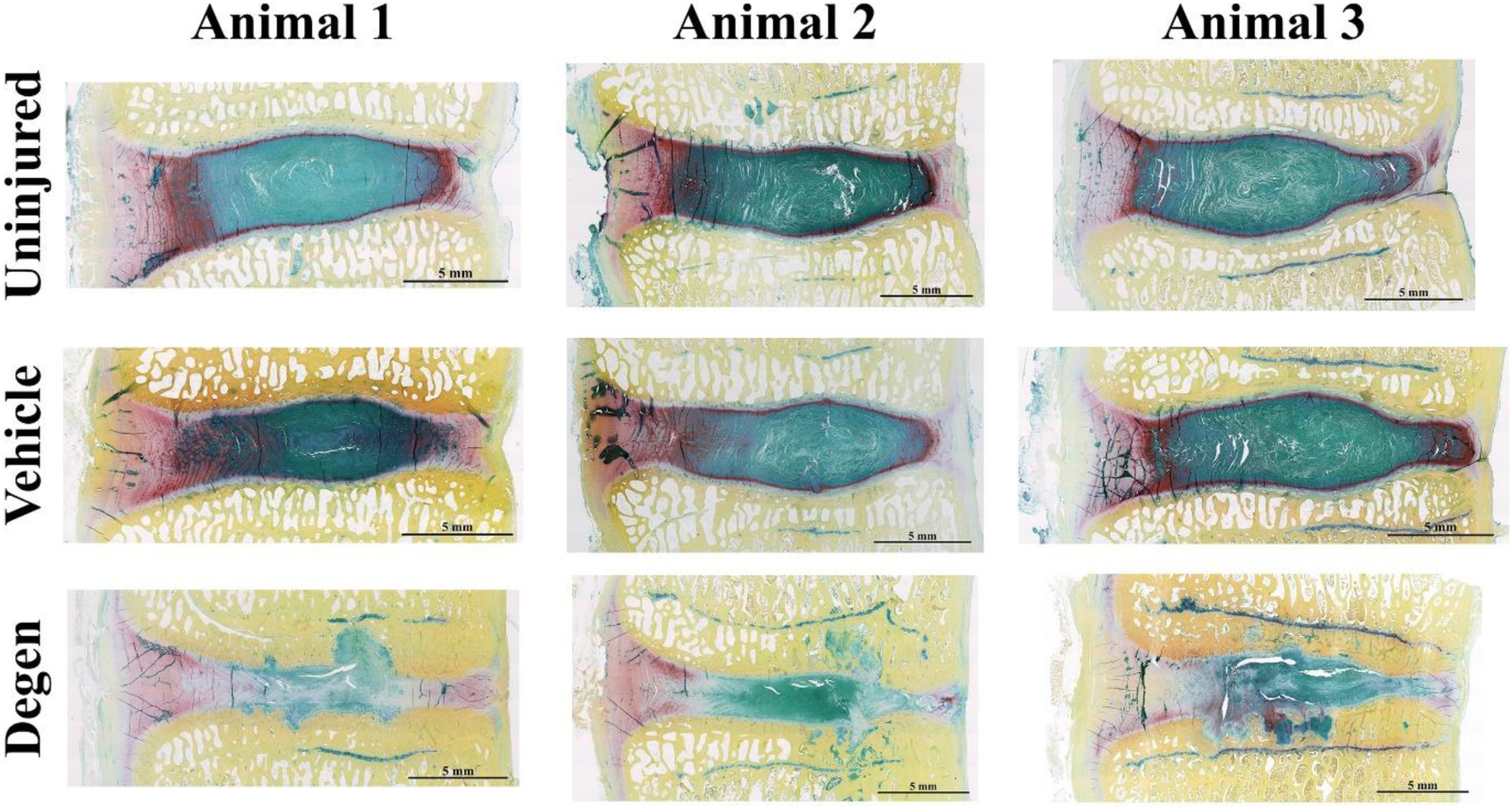
FAST-stained micrographs used for Rutges scoring. Mid-sagittal sections from Uninjured, Vehicle, and Degen IVDs from three respective animals were stained for histological analysis using the Rutges scoring system. Light Green: Outer AF; Dark Red: Inner AF; Teal: NP; Yellow: Vertebral Bone.

**Supplement 6.**
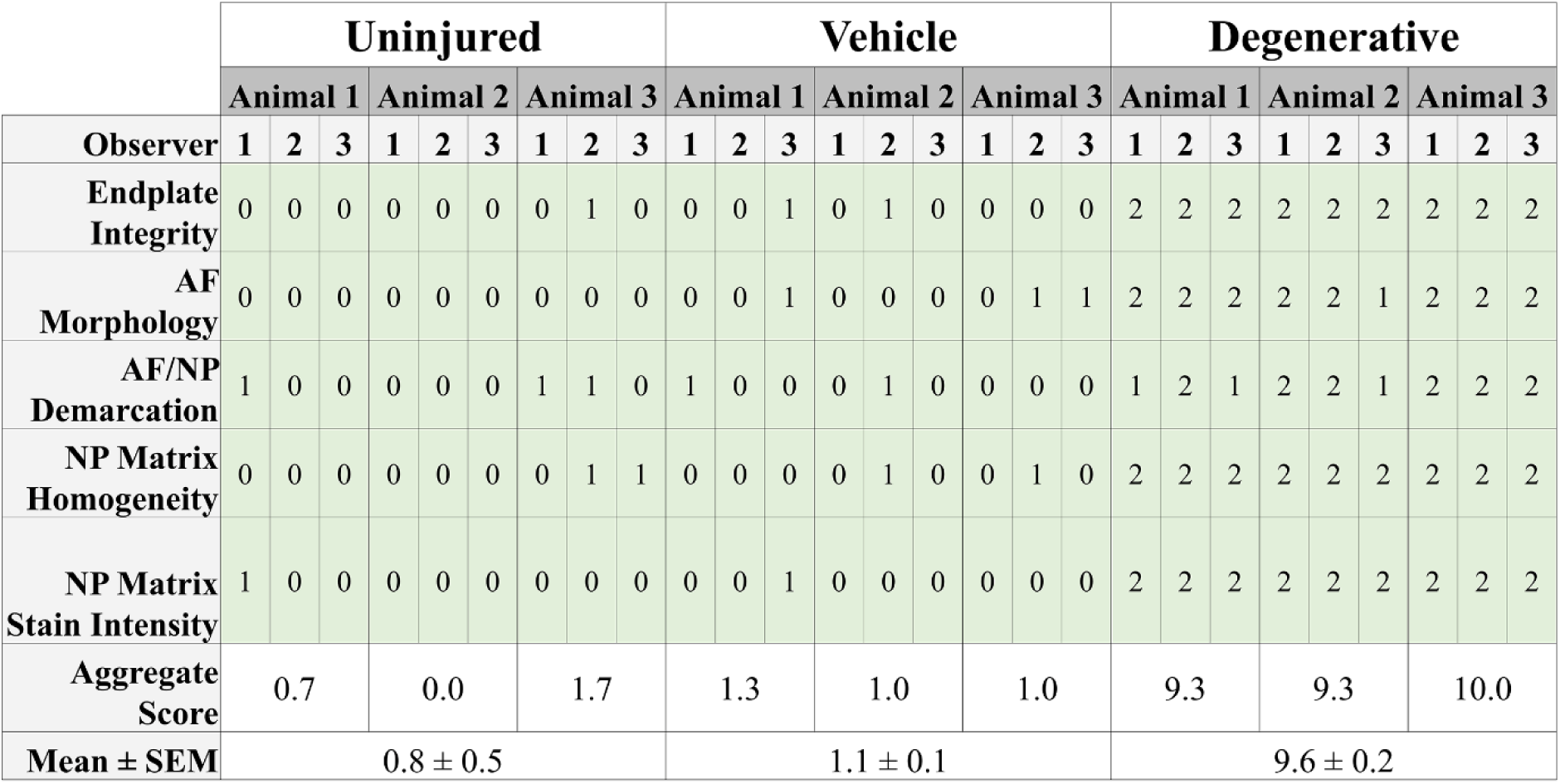
Raw Rutges scores for each animal and observer. Each sample was scored independently by two blinded and one non-blinded observers to determine the inter-observer reliability, which showed excellent agreement (ICC:0.94). Observers’ aggregate scores for each IVD were then averaged across observers for subsequent analysis. SEM: Standard Error of the Mean.

## References

1. Urban JPG, Roberts S. Degeneration of the intervertebral disc. Arthritis Res Ther. 2003;5:120.

2. Moon SM, Yoder JH, Wright AC, Smith LJ, Vresilovic EJ, Elliott DM. Evaluation of intervertebral disc cartilaginous endplate structure using magnetic resonance imaging. Eur Spine J. 2013;22:1820–8.

3. Katz JN. Lumbar disc disorders and low-back pain: socioeconomic factors and consequences. J Bone Joint Surg Am [Internet]. The Journal of Bone and Joint Surgery, Inc.; 2006 [cited 2015 Feb 9];88 Suppl 2:21–4. Available from: http://jbjs.org/content/88/suppl_2/21.abstract

4. Masuda K, Lotz JC. New Challenges for Intervertebral Disc Treatment Using Regenerative Medicine. Tissue Eng Part B Rev. 2010;16:147–58.

5. Surgeons AA of O. The Burden of Musculoskeletal Diseases in the United State: Prevalence, Societal, and Economic Cost. Second. Rosemont, IL: American Academy of Orthopaedic Surgeons; 2008.

6. Kushchayev S V., Glushko T, Jarraya M, Schuleri KH, Preul MC, Brooks ML, Teytelboym OM. ABCs of the degenerative spine. Insights Imaging. 2018;epub (ahea:1–22.

7. Wang S, Rui Y, Lu J, Wang C. Cell and molecular biology of intervertebral disc degeneration: current understanding and implications for potential therapeutic strategies. Cell Prolif. 2014;47:381–90.

8. Roberts S, Caterson B, Menage J, Evans EH, Jaffray DC, Eisenstein SM. Matrix metalloproteinases and aggrecanase: their role in disorders of the human intervertebral disc. Spine (Phila Pa 1976). 2000;25:3005–13.

9. Goupille P, Jayson MI, Valat JP, Freemont AJ. Matrix metalloproteinases: the clue to intervertebral disc degeneration? Spine (Phila Pa 1976). 1998;23:1612–26.

10. Johnson ZI, Schoepflin ZR, Choi H, Shapiro IM, Risbud M V. Disc in flames: Roles of TNF-α and IL-1β in intervertebral disc degeneration. Eur Cell Mater. 2015;30:104–16; discussion 116-7.

11. Inoue N, Espinoza Orías AA. Biomechanics of intervertebral disk degeneration. Orthop Clin North Am. 2011;42:487–99, vii.

12. Muriuki MG, Havey RM, Voronov LI, Carandang G, Zindrick MR, Lorenz MA, Lomasney L, Patwardhan AG. Effects of motion segment level, Pfirrmann intervertebral disc degeneration grade and gender on lumbar spine kinematics. J Orthop Res. 2016;34:1389–98.

13. Wilder DG, Pope MH, Frymoyer JW. The biomechanics of lumbar disc herniation and the effect of overload and instability. J Spinal Disord. 1988;1:16–32.

14. O’Connell GD, Leach JK, Klineberg EO. Tissue Engineering a Biological Repair Strategy for Lumbar Disc Herniation. Biores Open Access. 2015;4:431–45.

15. van Uden S, Silva-Correia J, Oliveira JM, Reis RL. Current strategies for treatment of intervertebral disc degeneration: Substitution and regeneration possibilities. Biomater Res. 2017.

16. Sloan SR, Lintz M, Hussain I, Hartl R, Bonassar LJ. Biologic Annulus Fibrosus Repair: A Review of Preclinical In Vivo Investigations. Tissue Eng Part B Rev. 2018;epub (ahea.

17. Reitmaier, Sandra, Graichen, Friedmar, Shirazi-Abd, Aboulfazl, Schmidt H. Seperate the sheep from the goats. J Bone Jt Surg. Elsevier B.V.; 2017;e102:1–11.

18. Alini M, Eisenstein SM, Ito K, Little C, Kettler AA, Masuda K, Melrose J, Ralphs J, Stokes I, Wilke HJ. Are animal models useful for studying human disc disorders/degeneration? Eur Spine J. 2008.

19. Hoogendoorn R, Doulabi BZ, Huang CL, Wuisman PI, Bank R a, Helder MN. Molecular changes in the degenerated goat intervertebral disc. Spine (Phila Pa 1976). 2008;33:1714–21.

20. Hoogendoorn RJW, Helder MN, Kroeze RJ, Bank R a, Smit TH, Wuisman PIJM. Reproducible long-term disc degeneration in a large animal model. Spine (Phila Pa 1976). 2008;33:949–54.

21. Hoogendoorn RJ, Wuisman PI, Smit TH, Everts VE, Helder MN. Experimental intervertebral disc degeneration induced by chondroitinase ABC in the goat. Spine (Phila Pa 1976). 2007;32:1816–25.

22. Gullbrand Y Z X SE, Malhotra NR, Schaer TP, Zawacki Z, Martin Y Z JT, Bendigo Y Z JR, Milby AH, Dodge GR, Vresilovic EJ, Elliott DM, Mauck RL, Smith LJ, Gullbrand SE, Malhotra NR, Schaer TP, Zawacki Z, Martin JT, Bendigo JR, Milby AH, Dodge GR, Vresilovic EJ, Elliott DM, Mauck RL, Smith LJ. A large animal model that recapitulates the spectrum of human intervertebral disc degeneration. Osteoarthr Cartil [Internet]. Elsevier Ltd; 2017;25:146–56. Available from: http://dx.doi.org/10.1016/j.joca.2016.08.006

23. Wilke HJ, Kettler A, Claes LE. Are sheep spines a valid biomechanical model for human spines? Spine (Phila Pa 1976). 1997;22:2365–74.

24. Reid JE, Meakin JR, Robins SP, Skakle JMS, Hukins DWL. Sheep lumbar intervertebral discs as models for human discs. Clin Biomech (Bristol, Avon). 2002;17:312–4.

25. Stewart DM, Monaco LA, Gregory DE. The aging disc: using an ovine model to examine age-related differences in the biomechanical properties of the intralamellar matrix of single lamellae. Eur Spine J. 2017;26:259–66.

26. Nisolle J, Bihin B, Kirschvink N, Neveu F, Clegg P, Dugdale A, Wang X, Vandeweerd J. Prevalence of Age-Related Changes in Ovine Lumbar Intervertebral Discs during Computed Tomography and Magnetic Resonance Imaging. Comp Med. 2016;66:300–7.

27. Hunter CJ, Matyas JR, Duncan NA. Cytomorphology of notochordal and chondrocytic cells from the nucleus pulposus: a species comparison. J Anat. 2004;205:357–62.

28. Reitmaier S, Kreja L, Gruchenberg K, Kanter B, Silva-Correia J, Oliveira JM, Reis RL, Perugini V, Santin M, Ignatius A, Wilke HJ. In vivo biofunctional evaluation of hydrogels for disc regeneration. Eur Spine J. 2014;23:19–26.

29. Sasaki M, Takahashi T, Miyahara K, Hirose aT. Effects of chondroitinase ABC on intradiscal pressure in sheep: an in vivo study. Spine (Phila Pa 1976). 2001;26:463–8.

30. Ghosh P, Moore R, Vernon-Roberts B, Goldschlager T, Pascoe D, Zannettino A, Gronthos S, Itescu S. Immunoselected STRO-3 ^+^ mesenchymal precursor cells and restoration of the extracellular matrix of degenerate intervertebral discs. J Neurosurg Spine [Internet]. 2012;16:479–88. Available from: http://thejns.org/doi/10.3171/2012.1.SPINE11852

31. Pfirrmann CW, Metzdorf A, Zanetti M, Hodler J, Boos N. Magnetic resonance classification of lumbar intervertebral disc degeneration. Spine (Phila Pa 1976). 2001;26:1873–8.

32. Borem R, Madeline A, Walters J, Mayo H, Gill S, Mercuri J. Angle-ply biomaterial scaffold for annulus fibrosus repair replicates native tissue mechanical properties, restores spinal kinematics, and supports cell viability. Acta Biomater. 2017;58.

33. O’Connell GD, Jacobs NT, Sen S, Vresilovic EJ, Elliott DM. Axial creep loading and unloaded recovery of the human intervertebral disc and the effect of degeneration. J Mech Behav Biomed Mater. Elsevier Ltd; 2011;4:933–42.

34. Farndale RW, Buttle DJ, Barrett AJ. Improved quantitation and discrimination of sulphated glycosaminoglycans by use of dimethylmethylene blue. BBA - Gen Subj. 1986;883:173–7.

35. Cissell DD, Link JM, Hu JC, Athanasiou KA. A Modified Hydroxyproline Assay Based on Hydrochloric Acid in Ehrlich’s Solution Accurately Measures Tissue Collagen Content. Tissue Eng Part C Methods. 2017;23:243–50.

36. Neuman RE, Logan MA. The determination of hydroxyproline. J Biol Chem. 1950;184:299–306.

37. Thompson JP, Pearce RH, Schechter MT, Adams ME, Tsang IK, Bishop PB. Preliminary evaluation of a scheme for grading the gross morphology of the human intervertebral disc. Spine (Phila Pa 1976). 1990;15:411–5.

38. Leung VYL, Chan WCW, Hung SC, Cheung KMC, Chan D. Matrix remodeling during intervertebral disc growth and degeneration detected by multichromatic fast staining. J Histochem Cytochem. 2009;57:249–56.

39. Preibisch S, Saalfeld S, Tomancak P. Globally optimal stitching of tiled 3D microscopic image acquisitions. Bioinformatics. 2009;25:1463–5.

40. Schindelin J, Arganda-Carreras I, Frise E, Kaynig V, Longair M, Pietzsch T, Preibisch S, Rueden C, Saalfeld S, Schmid B, Tinevez JY, White DJ, Hartenstein V, Eliceiri K, Tomancak P, Cardona A. Fiji: An open-source platform for biological-image analysis. Nat Methods. 2012;9:676–82.

41. Walter BA, Torre OM, Laudier D, Naidich TP, Hecht AC, Iatridis JC. Form and function of the intervertebral disc in health and disease: A morphological and stain comparison study. J Anat. 2015;227:707–16.

42. Motulsky HJ, Brown RE. Detecting outliers when fitting data with nonlinear regression - A new method based on robust nonlinear regression and the false discovery rate. BMC Bioinformatics. 2006;7:1–20.

43. McGraw, K., Wong S. Forming inferences about some intraclass correlation coefficients. Psychol Methods. 1996;1:30–46.

44. Bland JM, Altman DG. Statistical methods for assessing agreement between two methods of clinical measurement. Lancet (London, England). 1986;1:307–10.

45. Matsuyama Y, Chiba K, Iwata H, Seo T, Toyama Y. A multicenter, randomized, double-blind, dose-finding study of condoliase in patients with lumbar disc herniation. J Neurosurg Spine. 2018;28:499–511.

46. Marinelli NL, Haughton VM, Muñoz A, Anderson PA. T 2 relaxation times of intervertebral disc tissue correlated with water content and proteoglycan content. Spine (Phila Pa 1976). 2009;34:520–4.

47. Modic, M. T., Steinberg, P. M., Ross, J. S., Masaryk, T. J. & Carter JR. Degenerative disk disease: assessment of changes in vertebral body marrow with MR imaging. Radiology. 1988;166.

48. Fujiwara A, Lim TH, An HS, Tanaka N, Jeon CH, Andersson GB, Haughton VM. The effect of disc degeneration and facet joint osteoarthritis on the segmental flexibility of the lumbar spine. Spine (Phila Pa 1976). 2000;25:3036–44.

49. Kirkaldy-Willis WH, Farfan HF. Instability of the lumbar spine. Clin Orthop Relat Res. 1982;110–23.

50. Panjabi, M. Brown, M. Lindahl, S. Irstam, L. Hermens M. Intrinisc disc pressure as a measure of integrity of the lumbar spine. Spine J. 1988;13:913–7.

51. Adams MA, McNally DS, Dolan P. “Stress” distributions inside intervertebral discs. The effects of age and degeneration. J Bone Joint Surg Br. 1996;78:965–72.

52. Johannessen W, Elliott DM. Effects of degeneration on the biphasic material properties of human nucleus pulposus in confined compression. Spine (Phila Pa 1976). 2005;30:E724–9.

53. Johannessen W, Cloyd JM, O’Connell GD, Vresilovic EJ, Elliott DM. Trans-endplate nucleotomy increases deformation and creep response in axial loading. Ann Biomed Eng. 2006;34:687–96.

54. Mwale F, Roughley P, Antoniou J. Distinction between the extracellular matrix of the nucleus pulposus and hyaline cartilage: a requisite for tissue engineering of intervertebral disc. Eur Cell Mater. 2004;8:58–63; discussion 63-4.

55. Iatridis JC, Setton LA, Weidenbaum M, Mow VC. Alterations in the mechanical behavior of the human lumbar nucleus pulposus with degeneration and aging. J Orthop Res. 1997;15:318–22.

56. Buckwalter JA. Aging and degeneration of the human intervertebral disc. Spine (Phila Pa 1976). 1995;20:1307–14.

57. O’Connell GD, Malhotra NR, Vresilovic EJ, Elliott DM. The Effect of Nucleotomy and the Dependence of Degeneration of Human Intervertebral Disc Strain in Axial Compression. Spine (Phila Pa 1976). 2011;36:1765–71.

58. Boos N, Weissbach S, Rohrbach H, Weiler C, Spratt KF, Nerlich AG. Classification of Age-Related Changes in Lumbar Intervertebral Discs. Spine (Phila Pa 1976). 2002;27:2631–44.

59. Roberts S, Evans H, Trivedi J, Menage J. Histology and pathology of the human intervertebral disc. J Bone Joint Surg Am [Internet]. The Journal of Bone and Joint Surgery, Inc.; 2006 [cited 2015 Jun 20];88 Suppl 2:10–4. Available from: http://jbjs.org/content/88/suppl_2/10.abstract

60. Dudli S, Liebenberg E, Magnitsky S, Lu B, Lauricella M, Lotz JC. Modic type 1 change is an autoimmune response that requires a proinflammatory milieu provided by the “Modic disc.” Spine J. 2017;epub (ahea.

61. Mok FPS, Samartzis D, Ebhc D, Karppinen J, Luk KDK, Orth M, Orth F, Fong DYT, Cheung KMC, Orth F. ISSLS Prize Winner: Prevalence, Determinants, and Association of Schmorl Nodes of the Lumbar Spine With Disc Degeneration A Population-Based Study of 2449 Individuals. Spine (Phila Pa 1976). 2010;35:1944–52.

62. Mattei TA, Rehman AA. Schmorl’s nodes: current pathophysiological, diagnostic, and therapeutic paradigms. Neurosurg Rev. 2014;37:39–46.

63. Cheung KMC, Samartzis D, Karppinen J, Mok FPS, Ho DWH, Fong DYT, Luk KDK. Intervertebral disc degeneration: New insights based on “skipped” level disc pathology. Arthritis Rheum. 2010;62:2392–400.

64. Paul CPL, Smit TH, de Graaf M, Holewijn RM, Bisschop A, van de Ven PM, Mullender MG, Helder MN, Strijkers GJ. Quantitative MRI in early intervertebral disc degeneration: T1rho correlates better than T2 and ADC with biomechanics, histology and matrix content. PLoS One. 2018;13:e0191442.

